# A kinase-independent function of cyclin-dependent kinase 6 promotes outer radial glia expansion and neocortical folding

**DOI:** 10.1101/2022.05.24.493266

**Authors:** Lei Wang, Jun Young Park, Fengming Liu, Kris Olesen, Shirui Hou, Jamy C. Peng, Jordan Infield, Anna C. Levesque, Yong-Dong Wang, Hongjian Jin, Yiping Fan, Jon P. Connelly, Shondra M. Pruett-Miller, Miaofen G. Hu, Philip W. Hinds, Young-Goo Han

**Affiliations:** Department of Developmental Neurobiology, St. Jude Children’s Research Hospital, 262 Danny Thomas Place, Memphis, TN 38105, USA; Department of Cell and Molecular Biology, St. Jude Children’s Research Hospital, 262 Danny Thomas Place, Memphis, TN 38105, USA; Center for Applied Bioinformatics, St. Jude Children’s Research Hospital, 262 Danny Thomas Place, Memphis, TN 38105, USA; Center for Advanced Genome Engineering, St. Jude Children’s Research Hospital, 262 Danny Thomas Place, Memphis, TN 38105, USA; Department of Medicine, Division of Hematology/Oncology, Tufts University School of Medicine, Boston, MA 02111, USA; Department of Developmental, Molecular and Chemical Biology, Tufts University School of Medicine, Boston, MA 02111, USA

**Keywords:** Neural progenitor, corticogenesis, microcephaly, neocortex, cyclin-dependent kinase 6

## Abstract

The neocortex, the center for higher brain function, first emerged in mammals and has become massively expanded and folded in humans, constituting almost half the volume of the human brain. Primary microcephaly, a developmental disorder in which the brain is smaller than normal at birth, mainly results from the number of neurons in the neocortex being reduced because of defects in neural progenitor cells (NPCs). Outer radial glia (oRGs), NPCs that are abundant in gyrencephalic species but rare in lisencephalic species, are thought to play key roles in the expansion and folding of the neocortex. However, how oRGs expand, whether they are necessary for neocortical folding, and whether defects in oRGs cause microcephaly remain important questions in the study of brain development, evolution, and disease. Here, we show that oRG expansion in mice, ferrets, and human cerebral organoids requires cyclin-dependent kinase 6 (CDK6), the mutation of which causes primary microcephaly via an unknown mechanism. In a mouse model in which increased Hedgehog signaling expands oRGs and intermediate progenitor cells and induces neocortical folding, CDK6 loss selectively decreased oRGs and abolished neocortical folding. Remarkably, this function of CDK6 in oRG expansion did not require its kinase activity, was not shared by the highly similar CDK4 and CDK2, and was disrupted by the mutation causing microcephaly. Therefore, our results indicate that CDK6 is conserved to promote oRG expansion; that oRGs are necessary for neocortical folding; and that defects in oRG expansion may cause primary microcephaly.

**Significance Statement:** Primary microcephaly, a disorder in which the brain is smaller than normal at birth, disproportionately affects the neocortex. Although outer radial glia (oRGs) expansion is hypothesized to be important in neocortical expansion and folding, it remains unknown whether oRGs are necessary for neocortical folding and whether defective oRGs cause microcephaly. Moreover, how oRGs expand is not well understood. A mutation in CDK6 causes microcephaly via an unknown mechanism. Here, we show that CDK6 promotes oRG expansion and neocortical folding. This function of CDK6 does not require its kinase activity but is disrupted by a mutation that causes microcephaly. Our findings show that CDK6 is conserved to expand oRGs and provide evidence that oRG defects disrupt neocortical growth and folding.

## Introduction

The neocortex, which is involved in higher-order sensory, motor, and cognitive functions, is a defining feature of mammalian brains. It first emerged in mammals and has expanded greatly to give rise to folded (gyrencephalic) brains in many mammalian lineages. Neocortical expansion and folding require two coordinated processes that depend on neural progenitor cells (NPCs), namely, the increased production of neural cells and their tangential dispersion (1–6). The primary NPCs are the ventricular radial glia (vRGs) (also called apical radial glia) that form the ventricular zone (VZ) at the apical side of the developing brain (7, 8). vRGs have a radial process that extends basally to the pial surface and serves as a scaffold for migrating neurons. vRGs generate neurons directly or through secondary progenitors, including intermediate progenitor cells (IPCs) and outer radial glia (oRGs), both of which delaminate from the ventricular surface and translocate basally to form the subventricular zone (SVZ) (9–14). Therefore, oRGs are also called basal radial glia, translocating radial glia, or intermediate radial glia. IPCs are apolar, produce neurons directly, and have a limited proliferation capacity. In contrast, oRGs are bipolar with a radial process, produce neurons directly or via IPCs, and have a prolonged proliferation capacity. These features enable oRGs to boost neuron production and tangential dispersion considerably. Accordingly, oRGs and SVZ are prominently expanded in gyrencephalic mammals, as compared with mammals with a small and smooth (lisencephalic) neocortex (15–17), suggesting that they play key roles in neocortical expansion and folding. Whether oRGs are necessary for neocortical expansion and folding and what mechanisms underlie oRG expansion are important questions with regard to understanding the development and evolution of mammalian brains.

Primary microcephaly or MCPH (which stands for “microcephaly primary hereditary”) is a developmental disorder characterized by the brain being smaller than normal, resulting in a head circumference that is more than three standard deviations below the mean for sex and ethnicity at birth. So far, 28 genes (designated MCPH1–28) have been identified as causative genes for MCPH (Online Mendelian Inheritance in Man^®^). These MCPH genes encode proteins that are involved in diverse cellular processes, including centriole biogenesis, mitotic spindle formation, microtubule dynamics, chromosomal condensation, DNA replication and repair, transcription, signaling, and ribosomal RNA processing (18–20). Mutations that disrupt these processes result in defective vRG proliferation, differentiation, and/or survival, ultimately leading to a reduced number of neurons and an abnormally small brain. Although the identification and characterization of MCPH genes have revealed important regulatory mechanisms for vRGs, it remains unknown whether MCPH genes affect oRGs or whether abnormal oRGs contribute to MCPH because of the paucity of a suitable model system. Of note, in cases of MCPH, the brain often shows a simplified neocortical gyration pattern (21), which is suggestive of defective oRGs.

We have previously shown that Hedgehog (HH) signaling is a conserved mechanism that is necessary and sufficient for oRG expansion (22, 23). HH signaling promotes the production of oRGs from vRGs and increases oRG self-renewal. Here, we show that HH signaling requires cyclin-dependent kinase 6 (CDK6) to expand oRGs. In primary microcephaly 12 (MCPH12), a missense mutation in the *CDK6* gene causes microcephaly with a simplified gyral pattern through unknown mechanisms (24). In a mouse model in which increased HH signaling expands oRGs and IPCs, leading to an expanded and folded neocortex, CDK6 loss selectively decreased the oRGs and abolished neocortical folding. CDK6 was also required for oRG expansion in ferrets and in human cerebral organoids. Mechanistically, CDK6 promoted the self-renewal of oRGs. Unexpectedly, CDK6 loss did not affect the cell cycle progression of oRGs and, consistent with this, its kinase activity was dispensable for oRG expansion; however, the MCPH12-causing mutation in *CDK6* disrupted the function of CDK6 in oRG expansion. Therefore, we reveal a conserved role for CDK6 in oRG expansion and provide evidence that oRGs are necessary for neocortical folding and that defective oRG expansion may cause MCPH.

## Results

### HH signaling requires CDK6 to induce neocortical folding

We have previously reported that increased HH signaling in vRGs and their progeny in *GFAP::Cre; SmoM2^fl/+^* (hereafter *SmoM2* mutant) mice expands oRGs and IPCs, leading to neocortical expansion and folding (22). SMOM2 is a mutant Smoothened protein that constitutively activates HH signaling. We have also shown that SMOM2 requires the downstream GLI2 transcription factor to induce cortical expansion and folding (22), suggesting that SMOM2 induces these changes by regulating transcription. Therefore, to understand the underlying molecular mechanisms, we performed RNA sequencing (RNA-seq) in wildtype and *SmoM2* mutant embryonic cortices at embryonic day 14.5 (E14.5) and identified differentially expressed genes.

The *Cdk6* gene was among the genes whose expression was significantly increased in *SmoM2* mutants (*SI Appendix*, Fig. S1A). Notably, *CDK6* is mutated in MCPH12 in humans (24), whereas *Cdk6* loss does not cause microcephaly in mice (25, 26). These differing phenotypes suggest that CDK6 plays a role in neurodevelopmental processes that are prominent in humans but absent from or less prominent in mice. *SmoM2* mutant mice display neocortical folding and expansion of oRGs and IPCs, which are key features of developing human brains but are not prominent or absent in wildtype mice. Therefore, to investigate the role of CDK6 during neocortical development mimicking that of humans, we removed CDK6 from *SmoM2* mutants (*GFAP::Cre; SmoM2^fl/+^; Cdk6*^−/−^, hereafter *SmoM2; Cdk6KO*). The CDK6 loss slightly, but not significantly, decreased the cortical volume in *SmoM2; Cdk6KO* mutants as compared with *SmoM2* mutants; however, it completely abolished SMOM2-induced cortical folding (Fig. 1*A*–*C*). Consistent with the decreased cortical volume and the loss of folding, CDK6 loss decreased the number of SATB2+ upper-layer neurons (Fig. 1*F* and *G*), which are selectively expanded in *SmoM2* mutants (22). We concluded that CDK6 is required for SMOM2 to expand upper-layer neurons and to induce neocortical folding.

**Fig 1.**
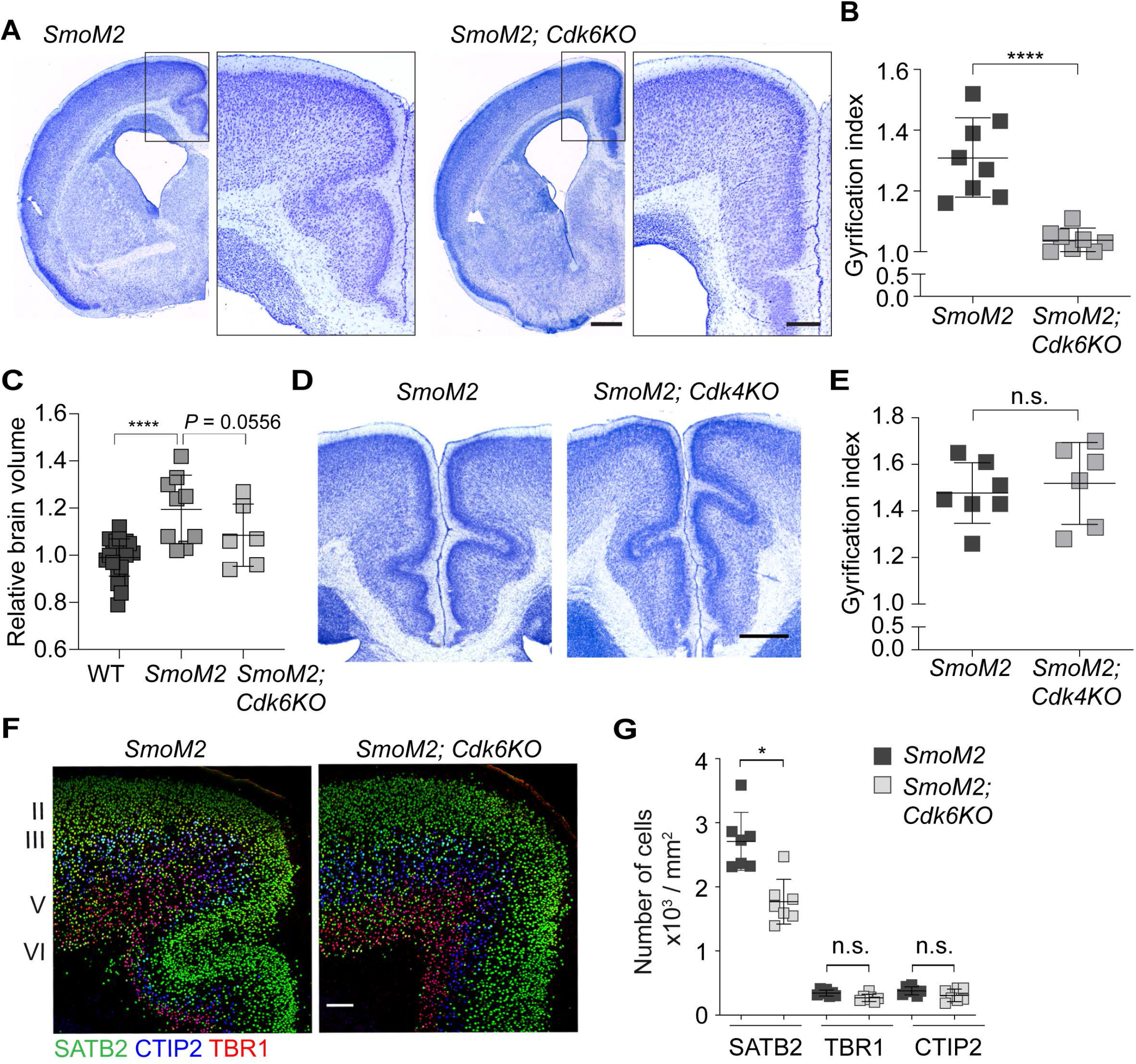
Loss of *Cdk6* but not of *Cdk4* abolished folding in the *SmoM2* mutant cortex. (*A*) Nissl-stained brain sections of P7 *SmoM2* and *SmoM2; Cdk6KO* mutants. Scale bar = 0.5 mm. Boxes show areas corresponding to the cingulate cortex that are magnified on the right. Scale bar = 0.2 mm. (*B*) Gyrification indices of the cingulate cortices at P7 (mean ± SD). **** *P* = 0.0002 (Mann–Whitney test; n = 8). (*C*) Relative brain volumes measured by MRI at P7 (mean ± SD). **** *P* < 0.0001 (Mann–Whitney test; n = 23 for control, 9 for *SmoM2*, 6 for *SmoM2; Cdk6KO*). (*D*) Nissl-stained brain sections of P7 *SmoM2* and *SmoM2; Cdk4KO* mutants. Scale bar = 0.5 mm. (*E*) Gyrification indices of the cingulate cortices at P7 (mean ± SD). n.s., *P* > 0.05 (Mann– Whitney test; n = 7 for *SmoM2*, 6 for *SmoM2; Cdk4KO*). (*F*) Micrographs of P7 *SmoM2* and *SmoM2; Cdk6KO* mutant brain sections immunolabeled for SATB2, CTIP2, and TBR1. Scale bar = 100 µm. (*G*) Quantification of the densities of neurons positive for SATB2, CTIP2, and TBR1 in the cingulate cortex at P7. * *P* < 0.05; n.s., *P* > 0.05 (Mann–Whitney test; n = 7 brain slices from three mice per group).

### CDK2 and CDK4 are dispensable for SMOM2-induced neocortical folding

CDK4 and CDK6 are highly similar kinases that are activated by the same D-type cyclins and play overlapping roles in G1 progression and G1/S transition during the cell cycle. Our data indicate that CDK4 cannot compensate for the loss of CDK6 in *SmoM2; Cdk6KO* mutants. Interestingly, our RNA-seq data showed that *Cdk4* mRNA levels were much higher than *Cdk6* mRNA levels in both control and *SmoM2* mutant embryos (*SI Appendix*, Fig. S1B). To investigate the role of CDK4 in SMOM2-induced neocortical expansion and folding, we removed CDK4 from *SmoM2* mutants (*GFAP::Cre; SmoM2^fl/+^; Cdk4*^−/−^, hereafter *SmoM2; Cdk4KO*). In contrast to CDK6, CDK4 was not required for *SmoM2* to induce neocortical folding (Fig. 1*D* and *E*). CDK2 is a kinase that regulates G1/S transition and S progression downstream of CDK4/6. Like CDK4, CDK2 was dispensable for SMOM2-induced neocortical folding (*SI Appendix*, Fig. S2). Together, these findings indicate that CDK6 plays unique and essential roles in neocortical folding in *SmoM2* mutants and that these roles are neither compensated for nor shared by CDK2 or CDK4.

### CDK6 loss selectively decreases oRGs

To investigate the mechanism by which CDK6 loss abolished cortical folding, we examined NPCs at E16.5, when mostly SATB2+ upper-layer neurons are produced. We previously showed that SMOM2 expands oRGs and IPCs (22). Remarkably, CDK6 loss selectively decreased the number of oRGs without affecting the numbers of vRGs and IPCs in *SmoM2; Cdk6KO* mutants (Fig. 2*A*, *C*, and *D*). Therefore, oRG expansion was necessary for neocortical folding and CDK6 was required for SMOM2 to expand oRGs. In contrast, CDK4 loss did not affect the number of oRGs, IPCs, or vRGs (Fig. 2*B*, *E*, and *F*), consistent with there being no change in neocortical folding in *SmoM2; Cdk4KO* mutants as compared with *SmoM2* mutants.

**Fig 2.**
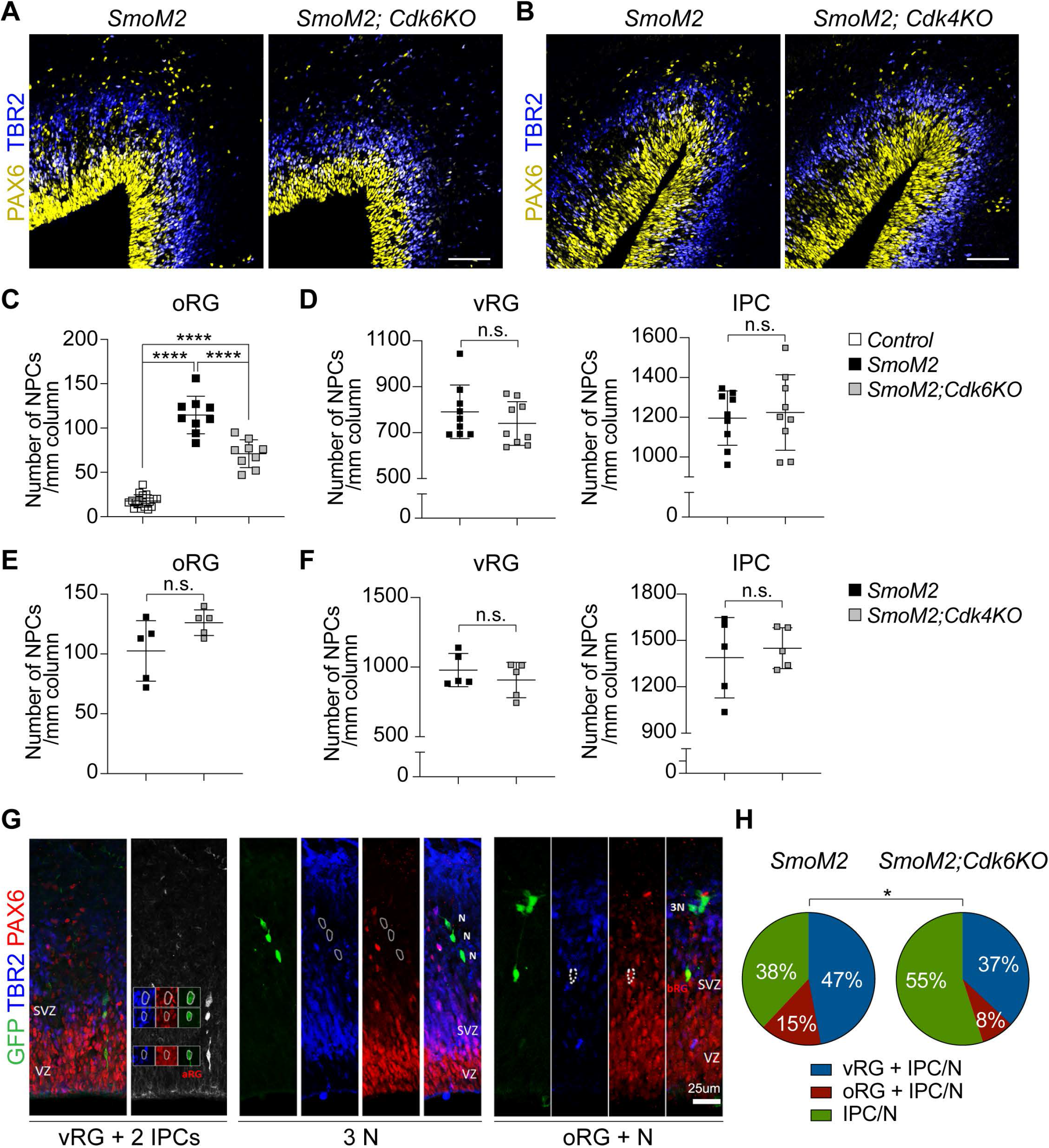
Loss of *Cdk6* but not of *Cdk4* decreased oRGs. (*A*, *B*) Micrographs of E16.5 cortices immunolabeled for PAX6 (yellow) and TBR2 (blue). Scale bar = 100 µm. (*C*) Quantification of oRG density in control, *SmoM2*, and *SmoM2;Cdk6KO* at E16.5 (mean ± SD). **** *P* < 0.0001 (unpaired *t*-test; n ≥ 9 brain slices from three mice per group). (*D*) Quantification of vRG and IPC densities in *SmoM2* and *SmoM2;Cdk6KO* at E16.5 (mean ± SD). n.s., *P* > 0.05 (unpaired *t*-test; n = 9 brain slices from three mice per group). (*E*, *F*) Quantification of oRG, vRG, and IPC densities in *SmoM2* and *SmoM2;Cdk4KO* at E16.5 (mean ± SD). n.s., *P* > 0.05 (unpaired *t*-test; n = 5 brain slices from three mice per group). (*G*) Micrographs showing examples of different types of clones. Cortices were labeled with GFP (green), PAX6 (red), and TBR2 (blue) to determine the cell fate 72 h (E16.5) after *in utero* intraventricular injection of GFP retroviruses at E13.5. Clones were defined as follows: IPC/N clones contained IPCs and/or neurons but no RGs; vRG+IPC/N clones had at least one vRG and IPC/N; oRG clones had at least one oRG and other cell types. (*H*) Distribution of clone types at E16.5. * *P* < 0.05 (chi-square test; 88 *SmoM2* clones, 67 *SmoM2;Cdk6KO* clones).

To confirm that SMOM2 required CDK6 to expand oRGs at the clonal level, we sparsely transduced vRGs with retroviruses expressing green fluorescent protein (GFP) at E13.5 and examined the fate of GFP+ cells in transduced clones containing at least two GFP+ cells at E16.5. In *SmoM2* mutants, 15% of clones contained oRGs with IPCs and/or neurons (Fig. 2*G* and *H*). In *SmoM2; Cdk6KO* mutants, clones containing oRGs decreased to almost half of that in *SmoM2* mutants (8%), similar to what was observed at the population level (Fig. 2*C*). Concomitantly, clones containing only IPCs and/or neurons increased from 38% in *SmoM2* mutants to 55% in *SmoM2; Cdk6KO* mutants (Figure 2*H*).

### SMOM2 requires CDK6 to promote oRG self-renewal

The best-known function of CDK6 is promoting cell cycle progression. Therefore, to investigate how CDK6 loss decreased the number of oRGs, we first examined the proliferation rate. We quantified the NPCs that incorporated 5-ethynyl-2′-deoxyuridine (EdU) 1.5 h after mice were injected with EdU at E16.5. CDK6 loss did not affect the proportions of oRGs, vRGs, and IPCs that were labeled by EdU (Fig. 3*A* and *B*). Because CDK6 is well known to promote G1/S transition, we tested whether the proportion of cells in G1 phase was affected by CDK6 loss. As reported previously (27), we identified G1-phase cells as those cells that expressed proliferating cell nuclear antigen (PCNA) but not geminin; PCNA is expressed in all proliferating cells, whereas geminin is present in all phases of the cell cycle except the G1 phase. The proportion of oRGs in the G1 phase was not affected by CDK6 loss (Figure 3*D* and *E*). Consistent with these findings, the proportion of oRGs expressing phosphorylated Retinoblastoma 1 (RB1), a CDK6 and CDK4 target that critically regulates G1/S transition and cell cycle progression, was not significantly affected by CDK6 loss (*SI Appendix*, Fig S3). Together, these findings suggest that CDK6 loss did not affect oRG proliferation. This is consistent with our previous findings that SMOM2 expands oRGs not by affecting their proliferation but by increasing the production of oRGs from vRGs and the self-renewal of oRGs (22). Therefore, we asked whether CDK6 loss affected the self-renewal of oRGs. We quantified the self-renewed fraction of oRGs remaining in the cell cycle 24 h after the previous S phase by injecting bromodeoxyuridine (BrdU) and EdU at 24 and 1.5 h, respectively, before collecting embryos at E16.5. The self-renewed fraction of oRGs (BrdU+ EdU+) was significantly smaller in *SmoM2; Cdk6KO* mutants than in *SmoM2* mutants (Fig. 3*A* and *C*), indicating that SMOM2 requires CDK6 to promote self-renewal of oRGs.

**Fig 3.**
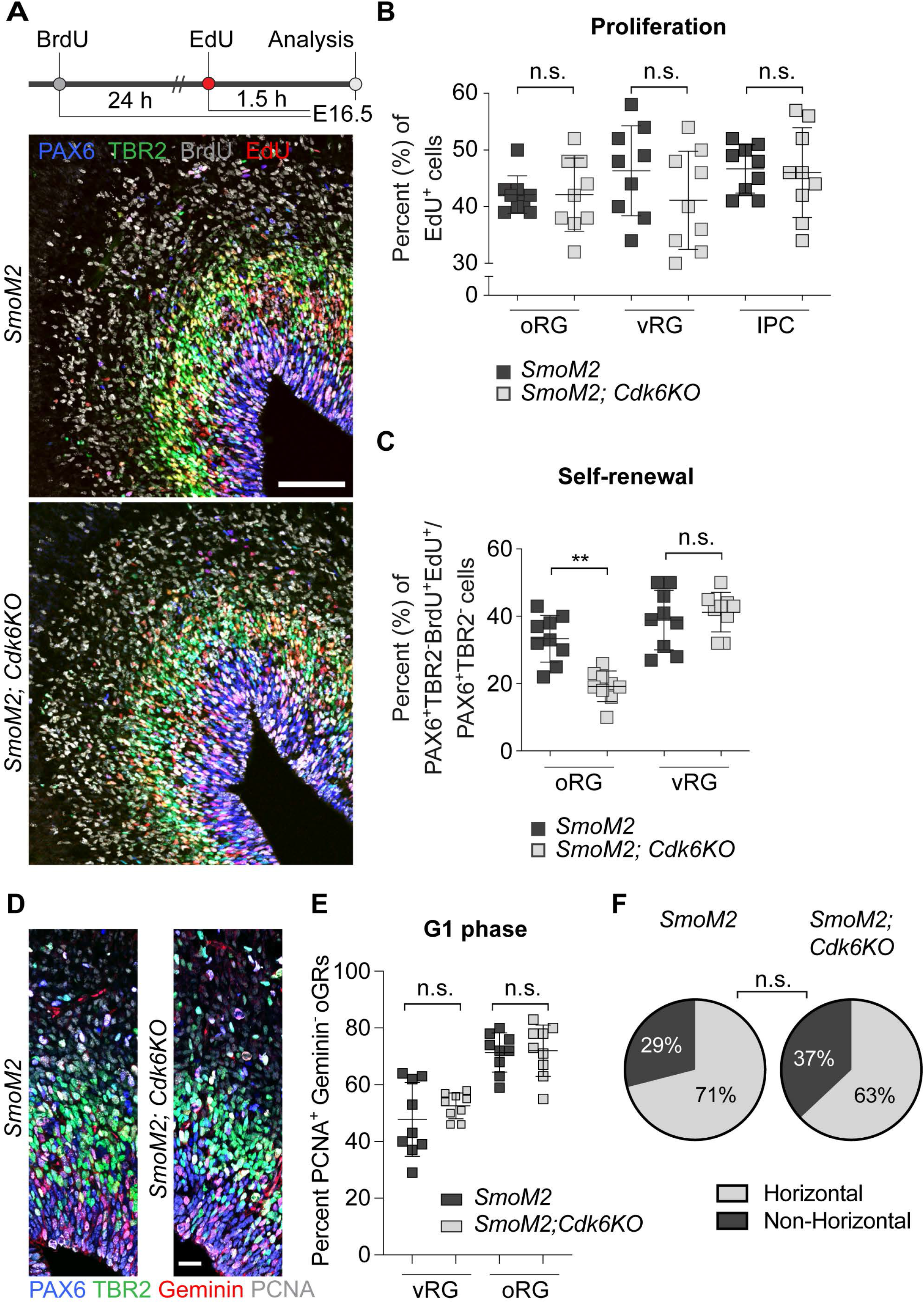
Loss of *Cdk6* decreased self-renewal of oRGs. (*A*) Schematic of experiments and micrographs of E16.5 cortices from embryos that were injected with BrdU (24 h before euthanasia) and EdU (1.5 h before euthanasia) and labeled for PAX6 (blue), TBR2 (green), BrdU (gray), and EdU (red). Scale bar = 100 µm. (*B*) Percentage of EdU+ oRGs, vRGs, and IPCs (mean ± SD). n.s., *P* > 0.05. (Mann–Whitney test; n = 9 brain slices from three mice per group) (*C*) Percentage of oRGs and vRGs that had self-renewed (PAX6^+^TBR2^−^BrdU^+^EdU^+^/PAX6^+^TBR2^−^ cells) (mean ± SD). ** *P* < 0.005; n.s., *P* > 0.05 (Mann–Whitney test; n = 9 brain slices from three mice per group). (*D*) Micrographs of E15.5 cortices labeled for PAX6 (blue), TBR2 (green), geminin (red), and PCNA (grey). Scale bar = 20 µm. (*E*) Quantification of the percentage of vRGs and oRGs in G1 phase (PAX6^+^ TBR2^−^ PCNA^+^ geminin^−^/PAX6^+^ TBR2^−^ PCNA^+^). n.s., *P* > 0.05 (Mann–Whitney test; n = 9 brain slices from three mice per group). (*F*) Quantifications of division modes of vRGs at E14.5. n.s., *P* > 0.05 (chi-square test; n = 27 for *SmoM2*, n = 75 for *SmoM2; Cdk6KO*).

Previous studies showed that vRGs dividing on an axis parallel to the ventricular surface (horizontal division) mostly produce neurons and IPCs, whereas vRGs dividing obliquely or vertically (non-horizontal division) produce oRGs (28, 29). SMOM2 promotes non-horizontal division of vRGs, increasing oRG production (22). Accordingly, we asked whether CDK6 affected the division angles of vRGs. The division angles of vRGs in *SmoM2* mutants and *SmoM2; Cdk6KO* mutants did not differ significantly (Fig. 3*F*), indicating that SMOM2 did not require CDK6 to increase non-horizontal division of vRGs for oRG production. A previous study showed that CDK6 localizes to the centrosomes in control fibroblasts but not in MCPH12 patient fibroblasts, which show supernumerary centrosomes and defective mitotic spindles, and that knockdown of CDK6 causes similar defects (24). However, we could not detect CDK6 in the centrosomes of vRGs and CDK6 loss did not result in supernumerary centrosomes or defective mitotic spindles. Taken together, our results suggest that CDK6 loss reduced the number of oRGs in *SmoM2; Cdk6KO* mutants by decreasing their self-renewal.

### Wildtype CDK6 but not MCPH12 mutant CDK6 expands oRGs

Our data show that CDK6 is necessary for SMOM2 to expand oRGs. We tested whether reintroducing CDK6 would restore oRGs in *SmoM2; Cdk6KO* mutants. Expression of human CDK6 in *SmoM2; Cdk6KO* mutants from E13.5 via *in utero* electroporation significantly increased the number of oRGs compared with electroporation of empty vectors expressing GFP only (Fig. 4*A* and *B*), confirming the role of CDK6 in SMOM2-driven oRG expansion. Reintroducing CDK6 also increased self-renewal of oRGs (*SI Appendix*, Fig. S4). A mutation changing the highly conserved alanine at position 197 of CDK6 to threonine (*CDK6^pA197T^*) was identified as a causal variant for MCPH12 (24). Remarkably, expression of CDK6^pA197T^ failed to increase oRGs in *SmoM2; Cdk6KO* mutants (Fig. 4*A* and *B*), suggesting that defective oRG expansion is a pathogenic mechanism for MCPH12. Next, we asked whether CDK6 was sufficient to expand oRGs. Expression of wildtype CDK6, but not of CDK6^pA197T^, in wildtype embryos at E13.5 significantly increased the number of oRGs by E16.5 (Fig. 4*C* and *D*), indicating that CDK6 is sufficient to expand oRGs in wildtype mice and that the MCPH12 mutation disrupts this function.

**Fig 4.**
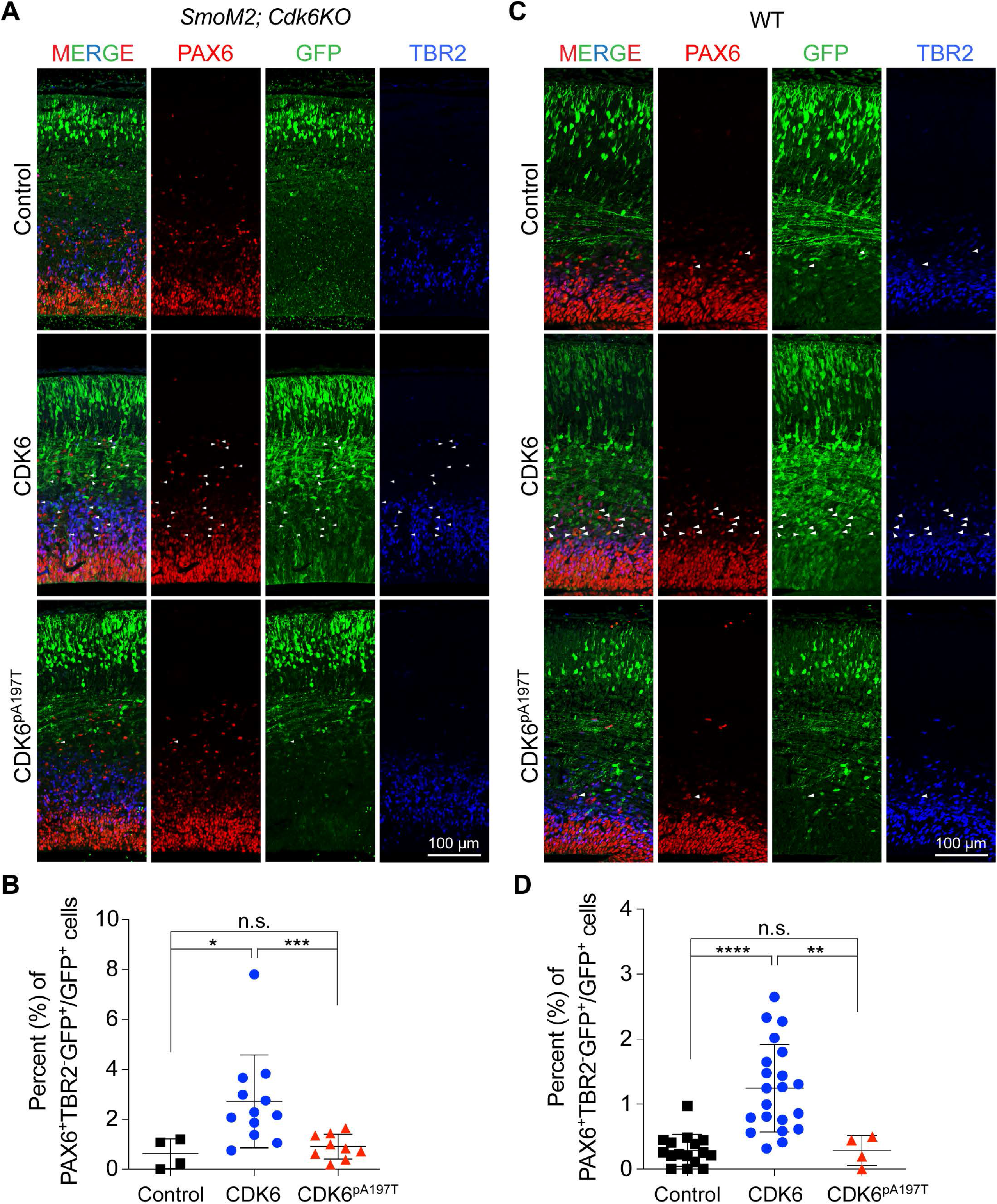
Wildtype CDK6, but not MCPH12 mutant CDK6 (CDK6^pA197T^), expanded oRGs. (*A*) Micrographs of E16.5 *SmoM2; Cdk6KO* brain sections stained for GFP (green), PAX6 (red), and TBR2 (blue) after *in utero* electroporation of plasmids expressing *GFP* alone or *GFP* with human *CDK6* or *CDK6^pA197T^*. Arrowheads indicate examples of GFP^+^ oRGs. (*B*) Percentage of oRGs among GFP+ cells (mean ± SD). *** *P* < 0.001; * *P* < 0.02; n.s., *P* > 0.05 (Mann–Whitney test; n = 4 brain slices from three embryos for controls, n = 12 brain slices from five embryos for *CDK6*, n = 9 brain slices from three embryos for *CDK6^pA197T^*). (*C*) Micrographs of E16.5 wildtype brain sections stained for GFP (green), PAX6 (red), and TBR2 (blue) after *in utero* electroporation of plasmids expressing *GFP* alone or *GFP* with *CDK6* or *CDK6^pA197T^*. Arrowheads indicate examples of GFP^+^ oRGs. (*D*) Percentage of oRGs among GFP+ cells (mean ± SD). **** *P* < 0.0001; ** *P* < 0.002; n.s., *P* > 0.05 (Mann–Whitney test; n = 16 brain slices from five embryos for controls, n = 21 brain slices from seven embryos for *CDK6*, n = 4 brain slices from two embryos for *CDK6^pA197T^*).

### The kinase function of CDK6 is dispensable for oRG expansion

Unexpectedly, CDK6, but not CDK4, was required for oRG expansion, independent of its well-known role in cell cycle regulation. To understand further the mechanism by which CDK6 expanded oRGs, we asked whether the kinase function of CDK6 was required for SMOM2 to expand oRGs. We crossed mice that have a point mutation disrupting their CDK6 kinase activity (*CDK6^pK43M^*) (30) with *SmoM2* mutants (*GFAP::Cre; SmoM2^fl/+^; Cdk6^pK43Mp/K43M^*, hereafter *SmoM2; CDK6^pK43M^*). In contrast to *SmoM2; CDK6KO* mutants, *SmoM2; CDK6^pK43M^* mutants developed neocortical folding (Fig. 5*A* and *B*). Moreover, CDK6^pA197T^ expressed in HEK293 cells was co-immunoprecipitated with cyclin D1 (Fig. 5*C*), an essential partner for its kinase activity, and phosphorylated RB1 in an *in vitro* kinase assay (Fig. 5*D* and *E*), suggesting that the pA197T mutation that causes microcephaly does not disrupt the kinase function of CDK6. Taken together, these findings suggest that CDK6 expands oRGs via a kinase-independent mechanism.

**Fig 5.**
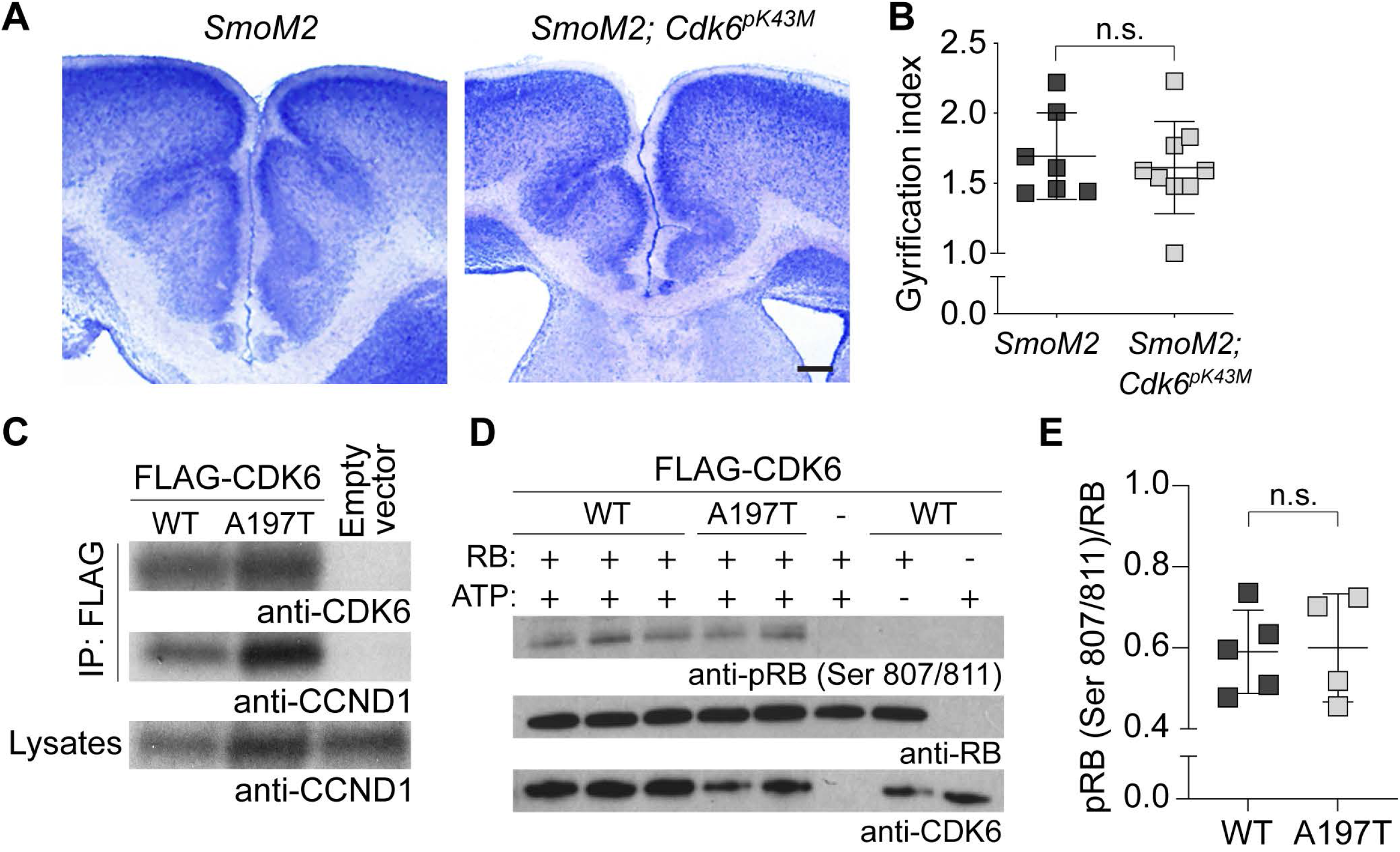
Kinase function of CDK6 was dispensable for oRG expansion. (*A*) Nissl-stained brain sections of P7 *SmoM2* and *SmoM2; Cdk6^K43M^* mutants. (*B*) Gyrification indices of the cingulate cortices at P7 (mean ± SD). n.s., *P* > 0.05 (Mann–Whitney test; n = 7 for *SmoM2*, n = 9 for *SmoM2; Cdk6^K43M^*). (C) Immunoblots, using antibodies to CDK6 and CCND1, of anti-FLAG antibody immunoprecipitants from HEK293 cells expressing FLAG-tagged CDK6 (WT), FLAG-tagged CDK6^pA197T^ (A197T), or no FLAG-tagged protein (empty vector). (*D*) Immunoblots for *in vitro* kinase assay of CDK6 and CDK6^pA197T^, using RB as a substrate. (*E*) Quantification of phospho-RB levels relative to total RB levels (mean ± SD). n.s., *P* > 0.05 (Mann–Whitney test; n = 5 for WT, n = 4 for A197T).

### CDK6 expands oRGs in ferrets

To investigate whether CDK6 expanded oRGs in a naturally gyrencephalic species, we cultured brain slices of E39 ferret embryos and, using a glass needle, locally infected the outer SVZ (OSVZ) with adenovirus expressing GFP alone or with human CDK6, shRNAs targeting *Cdk6*, or non-targeting shRNA (Fig. 6*A*). Adenoviruses expressing GFP alone or in conjunction with non-targeting shRNA were used as controls. Previous studies showed that adenovirus robustly transduced cortical progenitors, but not neurons, in embryonic ferret slices in culture (31, 32). In slices transduced with control virus, 45% of the GFP+ cells were oRGs by 3 days post infection (Fig. 6*B* and *C*). Knockdown of CDK6 significantly reduced the proportion of oRGs to 35%, whereas expression of CDK6 significantly increased this proportion to 76% (Fig. 6*B* and *C*). Of note, HH signaling activity that promotes oRG expansion is low in slices in culture (23), which could partly explain the relatively small effect of CDK6 knockdown. Taken together, our data suggest that CDK6 is necessary and sufficient to expand oRGs in ferrets.

**Fig 6.**
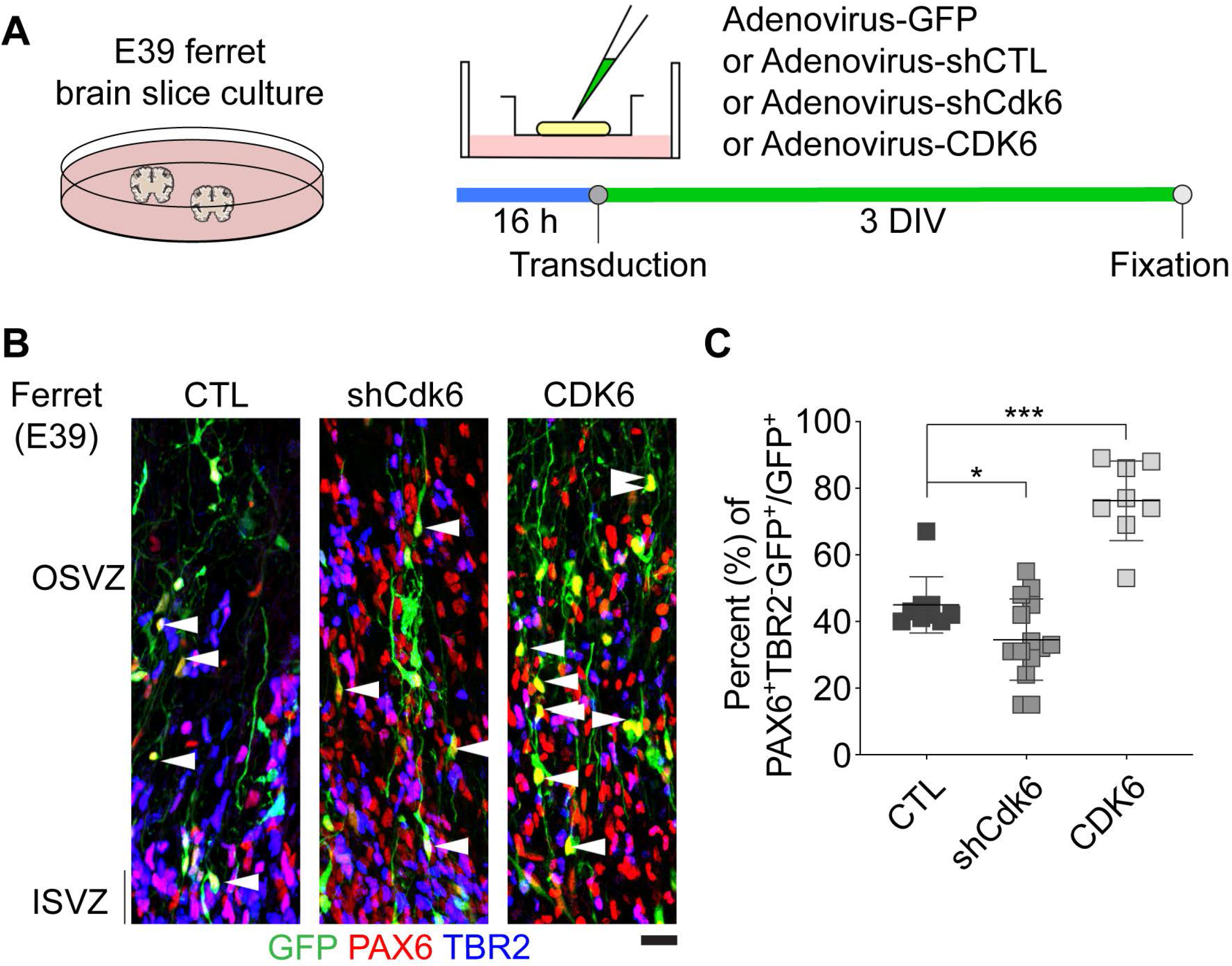
CDK6 expanded ferret oRGs. (*A*) Diagram of the experimental scheme. (*B*) Micrographs of E39 ferret brain slices transduced with adenoviruses expressing control vectors (shCTL or GFP), shCdk6, or human CDK6 and immunolabeled for PAX6 (red), TBR2 (blue), and GFP (green). Arrowheads indicate GFP^+^ PAX6^+^ TBR2^−^ oRGs in the OSVZ. Scale bar = 20 µm. (*C*) Percentages of transduced cells that remained as oRGs (GFP^+^ PAX6^+^ TBR2^−^/GFP^+^) 3 days after transduction (mean ± SD). *** *P* < 0.001; * *P* < 0.05 (unpaired *t*-test; n = 9 for control vectors, n = 14 for shCdk6, n = 8 for hCDK6 from two independent experiments).

### CDK6 is necessary for human oRG expansion in organoids

To investigate the role of CDK6 and the effects of an MCPH12 mutation (*CDK6^pA197T^*) in human oRGs by using a cerebral organoid model (33) (Fig. 7*A*), we generated *CDK6* knockout (*CDK6*^−/−^) and *CDK6^pA197T/pA197T^* (*CDK6^pA197T^*) knock-in human embryonic stem cells (ESCs). We generated two clonal cell lines for each genotype by using a CRISPR/Cas9 system (Fig. 7*B*). To examine alterations in potential off-target sites, we identified potential off-target sequences that match 12 bases upstream of PAM sequence and sequenced them from mutant ESCs. None of the potential off-target sequences of the gRNAs used was mutated in any cell line (not shown). All four mutant cell lines expressed pluripotency markers like the parental H9 human ESCs (*SI Appendix*, Fig. S5). We quantified oRGs in 7-week-old organoids generated from a wildtype and mutant hESCs. Both *CDK6* knockout and *CDK6^pA197T^* knock-in significantly decreased the number of oRGs in 7-week-old organoids (Fig. 7*C* and *D*) without affecting the division angles of the vRGs (Fig. 7*E* and *F*), which is consistent with the observation in *SmoM2; Cdk6KO* embryos. However, the change in the number of oRGs that were scarce compared to other cell types in organoids did not affect the overall size of organoids (not shown). Together, these findings suggest that CDK6 is required for human oRG expansion and that defective expansion of oRGs, a major NPC type in humans, is a pathogenic mechanism for microcephaly in patients with MCPH12.

**Fig 7.**
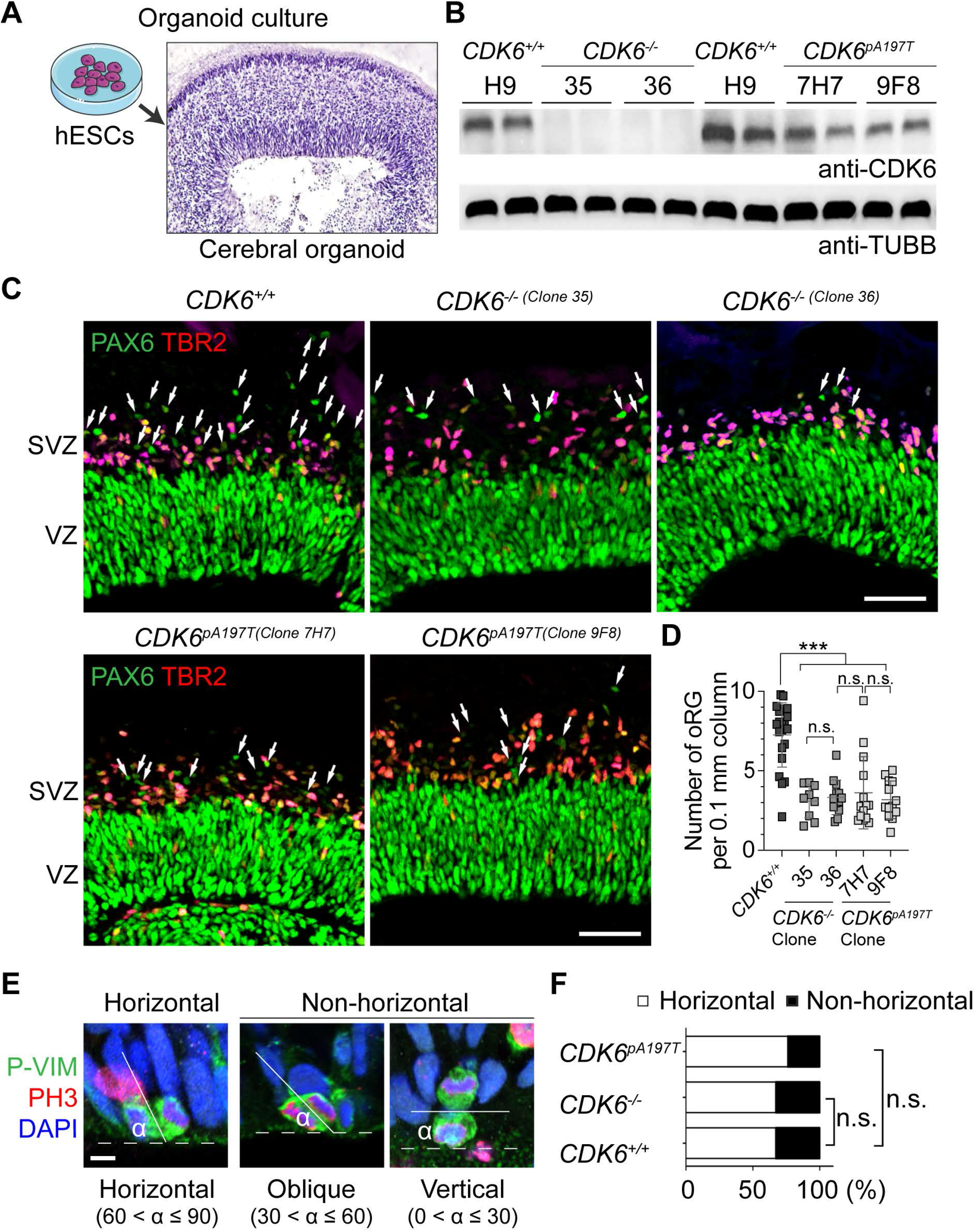
CDK6 was necessary for human oRG expansion in organoids. (*A*) An organoid section stained with hematoxylin, showing the VZ, SVZ, IZ, and CP. (*B*) Western blots of CDK6 from a parental H9 (*CDK6^+/+^*) and mutant ESC lines (*CDK6*^−/−^[clones 35 and 36] and *CDK6^pA197T^* [clones 9F8 and 7H7]). (*C*) Micrographs of cerebral organoid sections derived from the parental H9 and *CDK6* mutant ESC lines stained for PAX6 and TBR2. Scale bars = 50 µm. (*D*) Quantification of densities of oRGs. *** *P* < 0.001; n.s., *P* > 0.05 (Mann–Whitney test; 21 mini-cortices for H9 (*CDK6^+/+^*), 9, 13, 14, and 13 for clones 35, 36, 7H7, and 9F8, respectively). (*E*, *F*) Examples of horizontal, oblique, and vertical division of vRGs in cerebral organoids stained for phospho-vimentin (P-Vim), PH3, and DAPI and quantification of division modes of vRGs. n.s., *P* > 0.05 (Fisher exact test; 58 cells from 11 mini-cortices for *CDK6^+/+^*, 38 cells from 14 mini-cortices for *CDK6*^−/−^, 68 cells from 15 mini-cortices for *CDK6^A197T^*).

## Discussion

A missense mutation in *CDK6* was known to cause the microcephaly in MCPH12 (24); however, the pathogenic mechanism was unclear. Our study shows that CDK6 promotes oRG expansion in mice, ferrets, and human cerebral organoids and that the MCPH12 mutation disrupts this function of CDK6. MCPH disproportionately affects the neocortex, and patients frequently display a simplified neocortical gyrification pattern (18–21), signifying that the MCPH mutations primarily disrupt mechanisms responsible for neocortical expansion and folding. Studies have suggested that the expansion of oRGs and IPCs is critical for the expansion and folding of the neocortex (9, 10, 14–16, 34, 35). In particular, oRGs are greatly expanded in species with large/folded brains, especially humans, and oRG expansion is thought to underlie the complexity of the human brain (9, 10, 14, 36). Nevertheless, it was unknown whether oRGs were required for neocortical folding and whether MCPH mutations affected oRGs. We present evidence that oRGs are required for neocortical folding and that faulty oRG expansion can lead to MCPH.

vRGs have been shown to produce oRGs only within a restricted developmental window (23, 37). Beyond this window, oRGs have to be maintained for the remainder of the neurogenic period, which is protracted in species with large/folded brains. Therefore, genetic and environmental insults that disrupt oRG maintenance are likely to result in microcephaly. Our results show that CDK6 is required for oRG maintenance and that the MCPH12 mutation disrupts this function. Intriguingly, in ferrets, the expression levels of *Gli1*, a direct transcriptional target of HH signaling, and *Cdk6* are higher in the SVZ in prospective gyri (folds) than in prospective sulci (fissures) (38). Taken together with our results, these findings suggest that high HH signaling activity and consequent high CDK6 levels in prospective gyri expand and maintain oRGs, which in turn promote gyrus formation by increasing the local production and dispersion of neurons.

Unexpectedly, the kinase function of CDK6, which is essential for promoting cell cycle progression, was not required for oRG expansion. Consistent with this, CDK4, which plays largely redundant roles with CDK6 in cell cycle control, was not required for oRG expansion. How does CDK6 expand oRGs? Previous studies showed that CDK6, but not CDK4, interacts with transcription factors, including androgen receptor, RUNX1, STAT3, and C-JUN, to regulate transcription directly and that this transcriptional activity of CDK6 does not require kinase activity (39–41). In the hematopoietic system, these transcriptional activities inhibit the differentiation of progenitors and activate dormant hematopoietic stem cells and leukemic stem cells (39, 42). CDK6 also inhibits osteoclast and osteoblast differentiation by a mechanism unrelated to its cell cycle function (43, 44). In astrocytes, overexpression of CDK6, but not of CDK4, changes their shapes from multipolar to bipolar resembling progenitors and also alters their gene expression pattern toward that of immature glia (45). Similarly, CDK6 may regulate transcription to maintain oRGs in the progenitor state. Identifying CDK6-interacting proteins and assessing the transcriptional activities and targets of CDK6 in oRGs will help reveal molecular mechanisms for oRG expansion, neocortical development, and neurodevelopmental disorders.

## Materials and Methods

### Animals

All animal procedures were approved by the Institutional Animal Care and Use Committee of St. Jude Children’s Research Hospital.

The following mouse strains were used: *SmoM2^flox^* (The Jackson Laboratory [Jax], 005130), *GFAP::Cre (*JAX, 004600), *Cdk6^tm1Bbd^/Cnbc* (*Cdk6*^−/−^, EMMA, 06839), *Cdk4^tm2.1Bbd^/Cnbc* (*Cdk4*^−/−^, EMMA, 08029), *Cdk2*^−/−^ (ref. (46)), *Cdk6^pK43M^* (ref. (30)). Timed pregnant CD-1 mice were purchased from the Charles River Laboratories. Mice were maintained on a 12-h light/12-h dark cycle. We used mice of both sexes for experiments.

Timed pregnant ferrets (*Mustela putorius furo)* were purchased from Marshall BioResources and kept on a 12-h light/12-h dark cycle.

### Thymidine analogue injection

Solutions of 10 mg/mL BrdU (Sigma B-5002) and 2.5 mg/mL EdU (Invitrogen A10044) were prepared in sterile 0.9% NaCl solution. BrdU (50 μg/g) and EdU (10 μg/g) were injected intraperitoneally into the pregnant mice at 24 h and 1.5 h, respectively, before the animals were euthanized.

### Histologic processes

Mouse embryonic brains were dissected out, fixed overnight in 4% paraformaldehyde (PFA) at 4°C, cryoprotected for 24 h in 30% sucrose, embedded in OCT (Sakura Finetek), and sectioned at a thickness of 12 μm. Nissl staining was performed with cresyl violet (FD Neurotechnologies) according to the manufacturer’s instructions.

For immunolabeling, the cryosections were rinsed in PBS for 5 min and incubated with blocking solution (5% Normal Donkey Serum [Sigma-Millipore, D9663] and 0.1% Triton X-100 in PBS) for 1 h and primary antibodies in blocking solution overnight at 4°C. The slides were washed in PBS plus 0.1% Triton X-100 and incubated with secondary antibodies (1:400) in blocking solution for 2 h at RT. Coverslips were mounted with Aqua-Poly/Mount (Polysciences, Inc.). We used the following primary antibodies: anti-BrdU (Abcam, ab1893, 1:500), anti-CTIP2 (Abcam, ab18465, 1:500), anti-geminin (Santa Cruz Biotechnology, sc-13015, 1:50), anti-GFP (Novus Biologicals, NB100-1614, 1:500), anti-KI67 (DAKO, M7240, 1:100), anti-PAX6 (BioLegend, 901301, 1:300; R&D Systems, AF8150, 1:500), anti-PCNA (Santa Cruz Biotechnology, sc-56, 1:100); anti-phospho-histone H3 Ser10 (Abcam, ab14955, 1:1,000; Millipore-Sigma, 06-570, 1:2000), anti-pRB (Cell Signaling, D20B12, 1:500), anti-SATB2 (Millipore, ABE600, 1:2,000; Santa Cruz Biotechnology, sc-81376, 1:50), anti-TBR1 (Millipore, AB2261, 1:500), and anti-TBR2 (eBioscience, 14-4875-82, 1:250). We used Alexa Fluor®-conjugated secondary antibodies from Invitrogen. EdU staining was performed using a Click-iT EdU Kit (ThermoFisher, C10337) according to the manufacturer’s instructions.

### Microscopy and data analysis

Images were acquired on a Zeiss LSM 780 confocal microscope. Z-stacks of four or five optical sections at a step-size of 1 μm and a tiling of four to six images acquired using 20× objectives were combined for analysis. Quantification and analysis were carried out using ImageJ, Zen (Zeiss), and Photoshop (Adobe).

The VZ was defined as the area lining the ventricle and containing dense PAX6^+^ TBR2^−^ nuclei up to an area where cells uniformly expressed TBR2. The SVZ was defined as the cell-dense area containing uniformly TBR2^+^ cells above the VZ and below a cell-sparse area. vRGs were defined as PAX6^+^ TBR2^−^ cells in the VZ, and IPCs were defined as cells expressing TBR2 irrespective of PAX6 expression. oRGs were defined as PAX6^+^ TBR2^−^ cells present above the VZ. In organoids, the VZ was defined as the area lining the lumen and containing dense PAX6^+^ TBR2^−^ cells up to an area where the cells were sparser than in the VZ and formed stretches of TBR2^+^ cells. oRGs were quantified using criteria similar to those used for the mouse brain, i.e., PAX6^+^ TBR2^−^ cells above the VZ. A continuous line of TBR2^+^ cells formed the boundary between the VZ and SVZ. For analyses, only those VZ/SVZs that (1) were not obliquely cut, (2) were well separated from the neighboring VZ/SVZ, and (3) represented medial parts of the VZ/SVZ structures in serial sections were imaged.

To examine the division angles of vRGs, the cells were stained for P-Vim/PH3/DAPI. Only cells displaying clear separation of the chromosomes by cytoplasm, as revealed by P-Vim staining, were included in the analysis. The angles of division were measured using ImageJ.

### Plasmids

pBABE-GFP (Addgene 10668) and pLentiCRISPR-EGFP (Addgene 75159) were gifts from William Hahn and Beat Bornhauser (47), respectively. pCAG3-IRES-EGFP (pCIG3) was a gift from X. Cao (48). Wildtype *CDK6* and *CDK6^pA197T^* cDNA was cloned into pCIG3 by GenScript. FLAG-tagged wildtype *CDK6* and *CDK6^pA197T^* cDNA was cloned into in pcDNA3.1 by GenScript.

### Immunoprecipitation and Western blotting

Cells were lysed in modified RIPA buffer containing 20 mM Tris, pH 7.5, 150 mM NaCl, 0.5% Triton X-100, 1% SDS, 2mM EDTA, 10% glycerol, plus protease and phosphatase inhibitor cocktails (Sigma, PPC1010), passed through a 21-gauge needle three to five times, and centrifuged at 21,000 × *g* for 15 min at 4°C. The supernatants were incubated with primary antibodies and protein A/G agarose (Santa Cruz Biotechnology, sc-2003) for 2 h at 4°C. The protein samples were resolved by SDS-PAGE and transferred to PVDF membranes (Millipore). The membranes were blocked with 3% BSA + 0.05% Tween-20 for 1 h before incubation with primary antibodies overnight at 4°C. The following antibodies were used: anti-FLAG (Sigma-Aldrich, F1804), anti-CDK6 (Santa Cruz Biotechnology, sc-177; Cell signaling, 13331), and anti-cyclin D1 (Abcam, ab134175. 1:2000).

### *In vitro* kinase assay

*CDK6* KO ESCs were grown to 70% confluence then transfected with 2.5 µg of FLAG-WT CDK6 or FLAG-CDK6^pA197T^ plasmid per well by using Lipofectamine 3000. At 48 h after transfection, the growth medium was removed from the culture plates. The cells were rinsed briefly with ice-cold PBS and incubated on ice for 5 min in lysis buffer containing 20 mM Tris (pH 7.5), 150 mM NaCl, 1 mM EDTA, 1 mM EGTA, 1% Triton X-100, 2.5 mM Sodium pyrophosphate, 1 mM β-glycerophosphate, 1 mM Na_3_VO_4_, 1 μg/mL leupeptin (Cell Signaling, 9803s), and 1 mM PMSF. Samples were collected by scraping and transferred to microcentrifuge tubes on ice. Samples were centrifuged for 10 min at 14,000 × *g* at 4°C. The supernatants were transferred to new tubes, and the protein concentrations of the lysates were quantitated using a DC Protein Assay II Kit (BioRad, 5000112.). Protein A Dynabeads (ThermoFisher, 10001D) were briefly washed three times with lysis buffer. For each sample, 20 μL of beads were washed with PBS and incubated with 6 μg of anti-FLAG antibody (Sigma-Aldrich, F1804) in PBS + 0.5% BSA at 4°C for 3 h with gentle rocking. Beads were then washed three times with PBS and incubated with precleared lysate overnight at 4°C with gentle rocking. Lysates were precleared by incubating 250 μg of lysate, diluted to a volume of 200 μL, with 20 μL of washed beads for 1 h at 4°C with gentle rocking. After incubation with precleared lysates, beads were washed twice with lysis buffer and once with kinase buffer (25 mM Tris [pH 7.5], 2 mM DTT, 5 mM β-glycerophosphate, 0.1 mM Na_3_VO_4_, 10 mM MgCl_2)_ (Cell Signaling, 9802s) on ice. A quarter of the beads were then used for the *in vitro* kinase assay. Beads were incubated with kinase buffer containing 200 ng of human RB protein (Cell Signaling, 9303) and 200 µM ATP for 30 min at 37°C with gentle rocking. The reaction was terminated by adding Laemmli sample buffer to a concentration of 1×. Samples were then denatured at 100°C for 5 min and the beads were removed. Samples were separated by SDS-PAGE and transferred to PVDF membranes. Membranes were washed three times with TBS + 0.1% Tween-20 (TBST) for 10 min each, blocked with 5% BSA in TBST for 1 h, and incubated with primary antibody (diluted 1:1000 to 1:2000 in TBST + 5% BSA) overnight at 4°C with gentle rocking. Primary antibodies used for immunoblotting were anti-CDK6 (13331S), anti– phospho-RB (9308S), and anti-RB (9309S) (all from Cell Signaling). Membranes were washed with TBST, incubated for 90 min in HRP-linked secondary antibody (Jackson ImmunoResearch; 1:10,000 in TBST + 5% BSA), then washed three times with TBST and twice with TBS for 10 min each. Luminescent signal was induced using SuperSignal™ West Dura Extended Duration Substrate (ThermoScientific), and blots were exposed to film for 15 s to 2 min. The signal was analyzed using ImageJ.

### *In utero* intraventricular injection and clonal analysis

*In utero* viral injection and clonal analysis were performed as described previously (22). Replication-incompetent retroviruses were produced using HEK293T cell lines. Embryos were harvested 72 h (E16) post injection. Coronal sections of 50-μm thickness were collected and stained for GFP, PAX6, and TBR2. Clones that were clearly separated from each other and entirely included within the 50-μm thickness were imaged on the confocal microscope, using Z-stacks at step-intervals of 1 μm.

For *in utero* electroporation, we injected approximately 2 μL of DNA (2 μg/μL) mixed with 0.1% Fast Green (Sigma) in PBS into a lateral ventricle of the embryonic brain with a pulled glass micropipette. Square-wave electric pulses (30 V, 50 ms) were delivered five times at 950-ms intervals by using 3-mm tweezer-type electrodes and an electroporator (BTX ECM 830, Harvard Apparatus). Embryos were returned to their dam and allowed to develop until they were harvested for analysis.

### Magnetic resonance imaging (MRI)

MRI was performed as described previously (22).

### RNA isolation and sequencing

Total RNA was extracted from dissected cortices of E14.5 embryonic brains by using an RNeasy Micro Kit (Qiagen). A sequencing library was prepared using a TruSeq Stranded Total RNA Kit (Illumina) and sequenced with an Illumina HiSeq system. FASTQ sequences were mapped to the mm10 genome and counted with HTSEQ, and the Transcripts Per Kilobase Million (TPM) values were then computed. Statistical analyses and visualizations were performed using Partek Genomics Suite 6.6 and Stata MP/11.2 software.

### Ferret slice culture

E39 embryos were collected by caesarean section from pregnant ferrets anesthetized with isoflurane. After surgery, ferrets were euthanized with a barbiturate overdose. Organotypic slice cultures were established by published methods (32). E39 fetal brains were dissected out in ice-cold artificial cerebrospinal fluid (aCSF) [125 mM NaCl, 2.5 mM KCl, 1 mM MgCl_2_, 2 mM CaCl_2_, 1.25 mM NaH_2_PO_4_, 25 mM NaHCO_3_, 25 mM D-(+)-glucose], embedded in 4% low-melting-point agarose in aSCF without glucose, sectioned into 500-µm slices by using a Vibratome (Leica VT1200 S), transferred to culture inserts (MilliporeSigma™, PICM03050), and maintained in culture overnight in medium containing 66% BME (Gibco™, 21010046), 25% HBSS (Gibco™, 14170112), 5% (v/v) fetal bovine serum (FBS), 2 mM Glutamax (Gibco™, 35050061), 1% (v/v) N2 Supplement (Gibco™, 17502048), and 1% (v/v) penicillin–streptomycin at 37°C with 5% CO_2_. After the slices had been in culture for 16 h, adenoviruses (1 × 10^9^ CFU) (Vector Biolabs) were diluted in BME (1:10) and locally injected into the outer SVZ by using a FemtoJet® 4i microinjector (Eppendorf). After 72 h, the slices were fixed in 4% PFA for 1 h.

### Cerebral organoid culture

Organoids were grown in culture as described previously (22, 33). H9 human ESCs were obtained from the WiCell Research Institute and cultured in mTeSR1 medium (STEMCELL Technologies). Embryoid bodies (EBs) were grown by seeding 9000 cells in 150 μL of DMEM/F12 medium (Life Technologies, 11330-032) supplemented with 10% knockout serum replacement (Life Technologies, 10828-028), 3% FBS (Life Technologies, 10439-016), 1% GlutaMAX (Life Technologies, 35050-061), 1% MEM-NEAA (Life Technologies, 11140-050), 7 ppm (v/v) β-mercaptoethanol (Life Technologies, 21985-023), 4 ng/mL bFGF (Peprotech, 100-18B), and 50 μM Rho-associated kinase inhibitor (ATCC, ACS-3030) into each well of a 96-well Lipidure®-Coat Plate (Gel Company, LCV96). The medium was changed every other day for 6–7 days, omitting the bFGF and ROCK inhibitor after day 4. When the EBs attained a diameter of approximately 500 μm, they were transferred to wells of a Costar® 24-well plate (Corning 3473) (one or two EBs per well). The EBs were fed every other day with neural induction medium consisting of DMEM/F12 supplemented with 1% N2 supplement (Life Technologies, 17502-048), 1% GlutaMAX, 1% MEM-NEAA, and 1 μg/mL heparin for 4–5 days until neuroepithelial morphology became evident. The neuroepithelial aggregates were then embedded in a drop of Matrigel (Corning, 356234). The embedded aggregates (*n* = 16) were grown in 6-mm dishes containing 5 mL of differentiation medium (50% DMEM/F12, 50% Neurobasal Medium, 0.5% N2 supplement, 1% B27 without vitamin A [Life Technologies, 12587-010], 0.025% [v/v] human insulin [Sigma, I9278], 3.5 ppm [v/v] β-mercaptoethanol, 1% GlutaMAX, 0.5% MEM-NEAA, and 1% penicillin–streptomycin) with constant shaking at 75 rpm for 4 days, with the medium being changed on the second day. Four days after differentiation, the tissue droplets were fed with differentiation medium containing 1% B27 Supplement with vitamin A (Life Technologies 17504-044), instead of B27 without vitamin A, and incubated at 37°C in 5% CO_2_ with constant rotation at 75 rpm, with the medium being replenished every 3 days. From day 40, Matrigel was added to the differentiation medium at 1%.

### Cas9-CRISPR genome editing

To make *CDK6^-/-^* hESCs, guide RNAs targeting exon 3 (TTCAAACACTAAAGTTAGTT and TCAGTGGGCACTCCAGGCTC) of the *CDK6* locus were cloned into pLentiCRISPR-EGFP. Packaged lentiviral particles collected in mTeSR1 medium were added to fresh mTeSR1 in a 1:9 ratio (v/v) with 8 μg/mL polybrene and incubated overnight. One week after lentivirus transduction, H9 human ESCs were sorted into 96-well plates by GFP expression to make single-cell clones. Targeted deletions were screened by Sanger sequencing of purified PCR products amplified from genomic DNA.

To make *CDK6^pA197T^* hESCs, H9 hESCs were treated with StemFlex (Thermo Fisher Scientific) supplemented with 1X RevitaCell (Thermo Fisher Scientific) for 1 hour. H9 cells were nucleofected (Lonza, 4D-Nucleofector™ X-unit) with precomplexed ribonuclear proteins (RNPs) consisting of 250 pmol of chemically modified sgRNA (GUCCAGCUACGCCACCCCCG, Synthego), 165 pmol of Cas9 protein (St. Jude Protein Production Core), 500ng of pMaxGFP (Lonza), and 3ug of ssODN donor (ccattgcaggtcgtcacgctgtggtacagagcacccgaagtcttgctccagtccagctacAcAacccccgtggatctctggagtgttggctgcatatttgcagaaatgtttcgtagaaagtaagaaa, modification in upper case, IDT). The nucleofection was carried out according to the manufacturer’s recommended protocol. Five days post nucleofection, cells were sorted for single cells for transfected (GFP+) cells and plated into prewarmed (37C) StemFlex media supplemented with 1X CloneR (Stem Cell Technologies) onto Vitronectin XF (Stem Cell Technologies) coated plates. Clones were screened for the desired modification via targeted deep sequencing using gene specific primers with partial Illumina adapter overhangs on a Miseq Illumina sequencer as previously described (49). Clones containing the desired edit were identified, expanded, and sequence confirmed. Cell identity of the final clones was authenticated using the PowerPlex® Fusion System (Promega).

### Statistics

Statistical analysis was performed using the GraphPad Prism software. Data were pre-tested for normality by the Kolmogorov–Smirnov (K-S) test. For normal distribution data, an unpaired *t*-test was used. Non-parametric analysis methods, including the Mann– Whitney test, chi-square test, and Fisher exact test, were also used. *P* values of less than 0.05 were considered to indicate significance. No statistical methods were used to predetermine the sample size, but our sample sizes were similar to those generally employed in the field.

## Acknowledgments

We thank the staff of the Animal Resource Center, the Small Animal Imaging Center, the Hartwell Center for Bioinformatics and Biotechnology, and the Cell and Tissue Imaging Center at St. Jude Children’s Research Hospital for technical assistance. We thank David Finkelstein for help with the RNA-seq analyses and Xinwei Cao for the pCIG3 vector and for reviewing the manuscript. We thank Keith A. Laycock, PhD, ELS, for scientific editing of the manuscript. J.C.P. is supported by the American Lebanese Syrian Associated Charities (ALSAC). This work was supported by a Whitehall Foundation Research Grant, by the American Lebanese Syrian Associated Charities (ALSAC), and by the National Institutes of Health (R01NS100939 to Y.-G. H.). The content is solely the responsibility of the authors and does not necessarily represent the official views of the National Institutes of Health.

---

**Fig S1.**
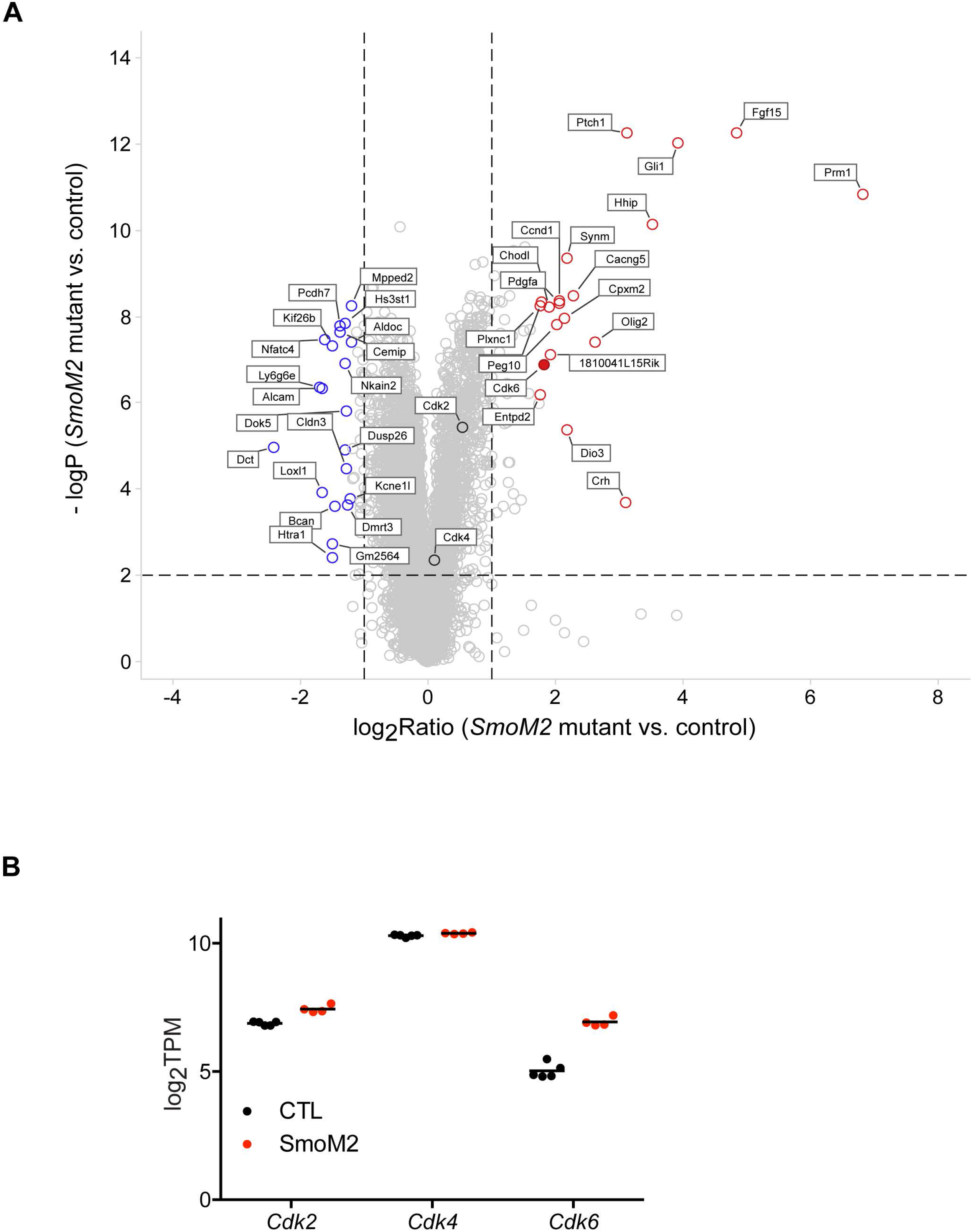
CDK6 was significantly increased in *SmoM2* mutant cortices. (A) Volcano plots showing gene expression ratios between control and *SmoM2* mutants and *P* values for differential expression obtained by RNA-seq from five control and four *SmoM2* mutant embryos. The top 20 genes with increased expression in *SmoM2* mutants are highlighted with red open circles, and the top 20 genes with decreased expression are highlighted with blue open circles. *Cdk6* is highlighted by a red filled circle. *Cdk4* and *Cdk2* are indicated by black open circles. (B) A plot showing the expression levels (log_2_TPM) of *Cdk2*, *Cdk4*, and *Cdk6* (mean ± SD).

**Fig S2.**
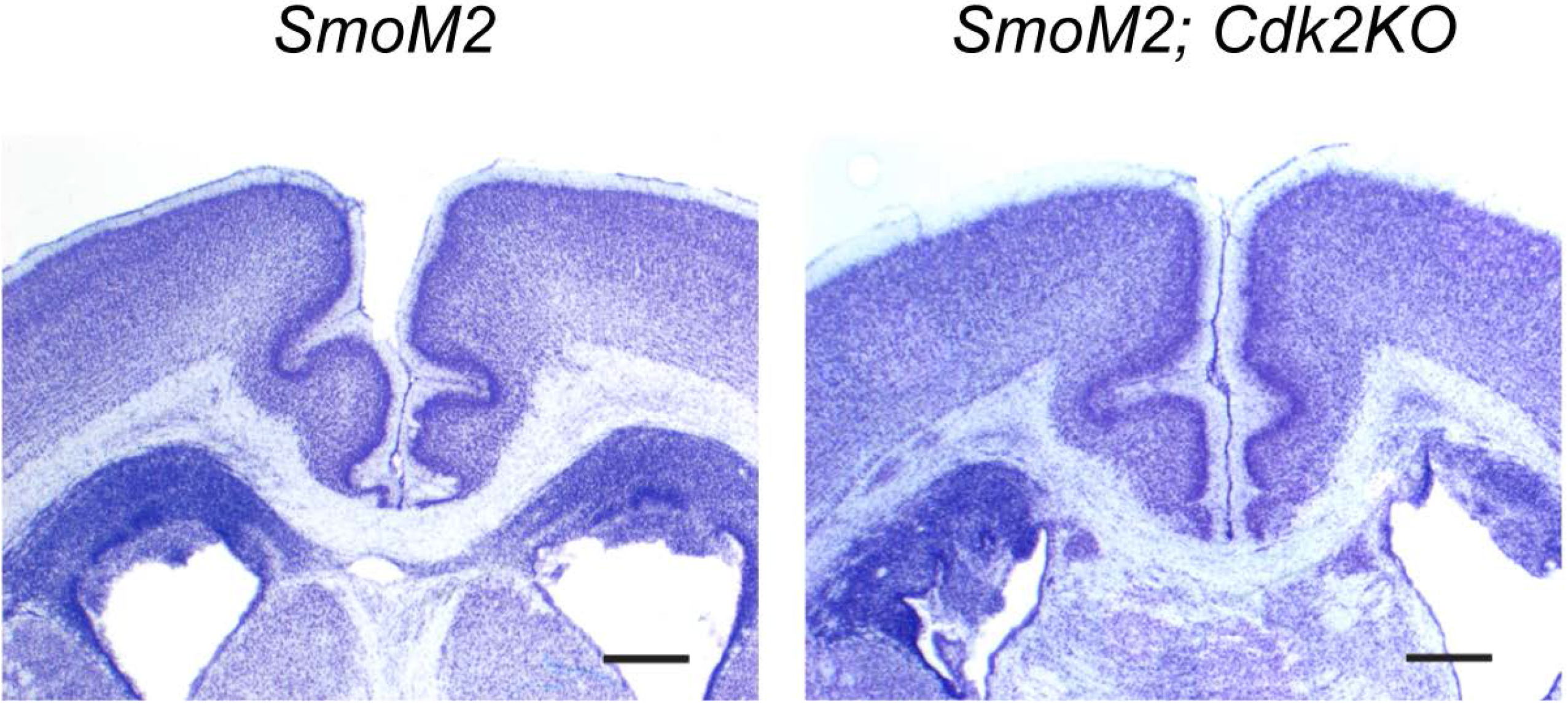
Loss of *Cdk2* did not affect *SmoM2*-induced neocortical folding. Nissl-stained brain sections of *SmoM2; Cdk2KO* mutants. Scale bar = 0.1 mm.

**Fig S3.**
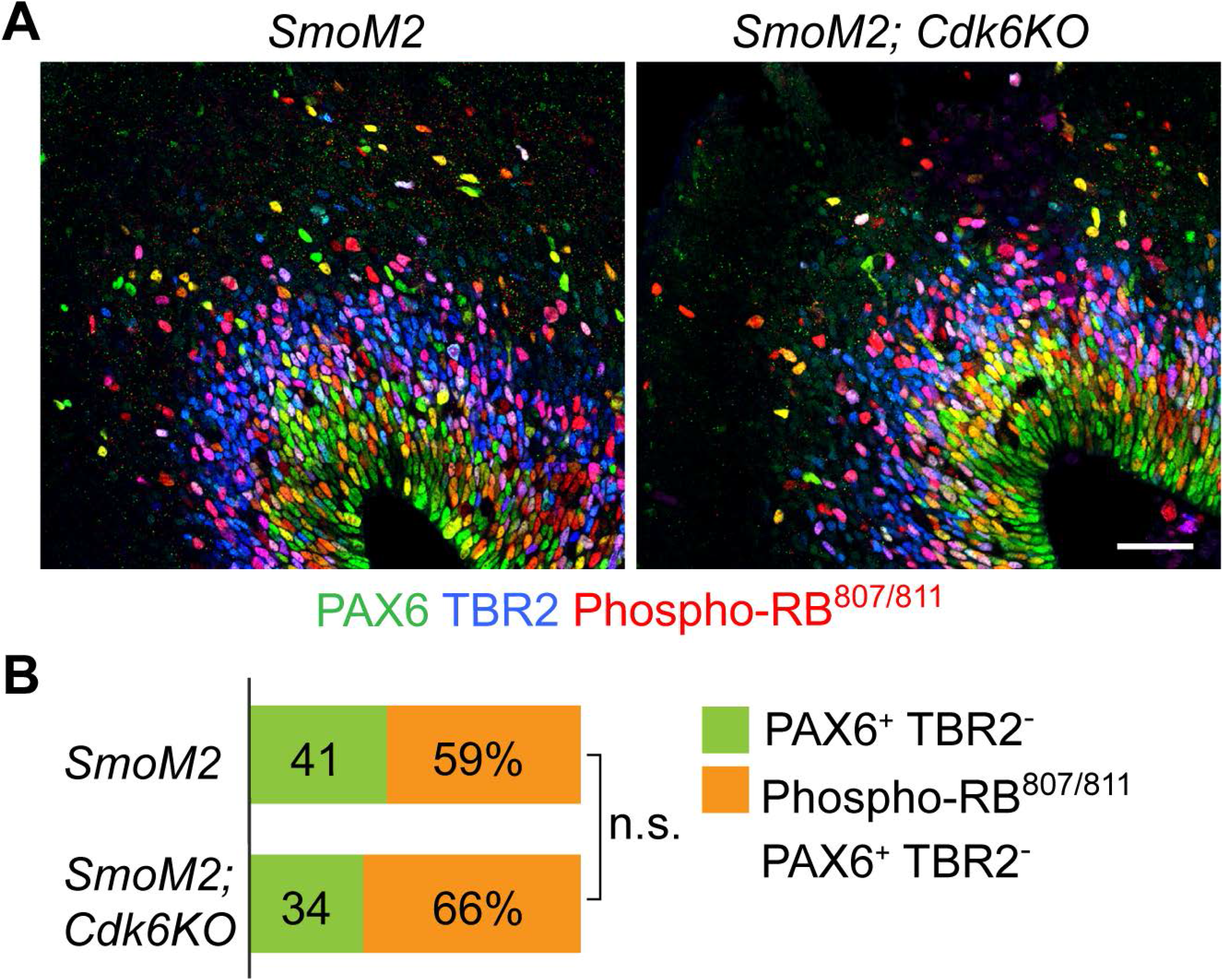
Loss of *Cdk6* did not affect the phosphorylation of RB^807/811^ in oRGs. (A) Micrographs of E16.5 cortices labeled for PAX6 (green), TBR2 (blue), and phospho-RB^807/811^ (red). Scale bar = 50 µm. (B) Percentage of oRG cells expressing phospho-RB^807/811^ (PAX6^+^ TBR2^+^ phospho-RB^807/811^). n.s., *P* > 0.05 (chi-square test; n = 3 embryos per group).

**Fig S4.**
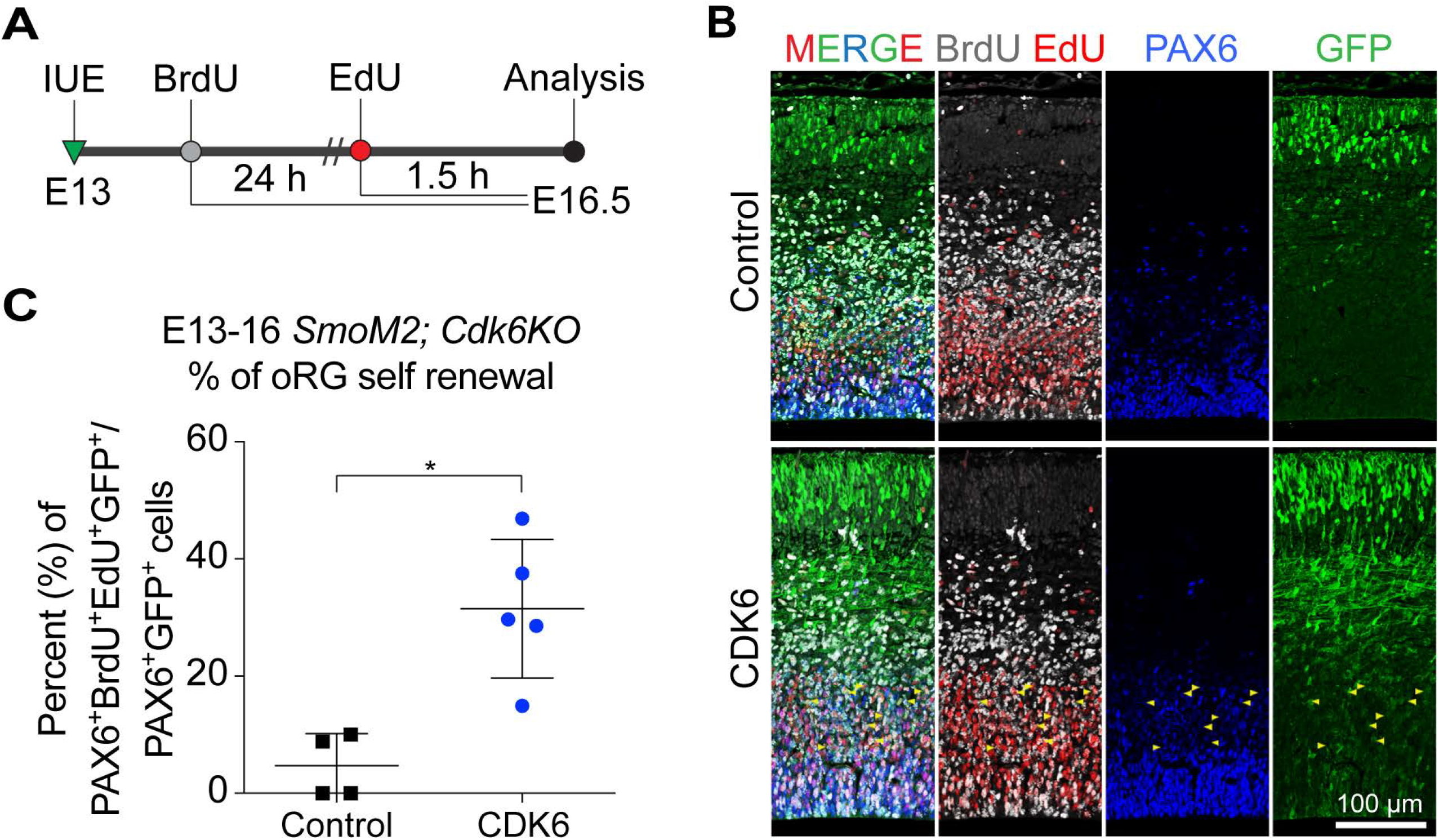
Electroporation of CDK6 rescued self-renewal of oRGs in *SmoM2; Cdk6KO*. (*A*) Diagram of the experimental scheme. (*B*) Micrographs of E16.5 brain sections from embryos that were electroporated with plasmids expressing GFP alone or with *CDK6* at E13.5 and injected with BrdU (24 h before euthanasia) and EdU (1.5 h before euthanasia). Sections were stained for PAX6 (blue), GFP (green), BrdU (gray), and EdU (red). Arrowheads indicate examples of PAX6^+^ BrdU^+^ EdU^+^ GFP^+^ oRGs. (C) Percentage of self-renewed oRGs among GFP-expressing oRGs (mean ± SD). * *P* < 0.02 (Mann–Whitney test; n = 4 brain slices from two embryos for controls, n = 5 brain slices from two embryos for CDK6).

**Fig S5.**
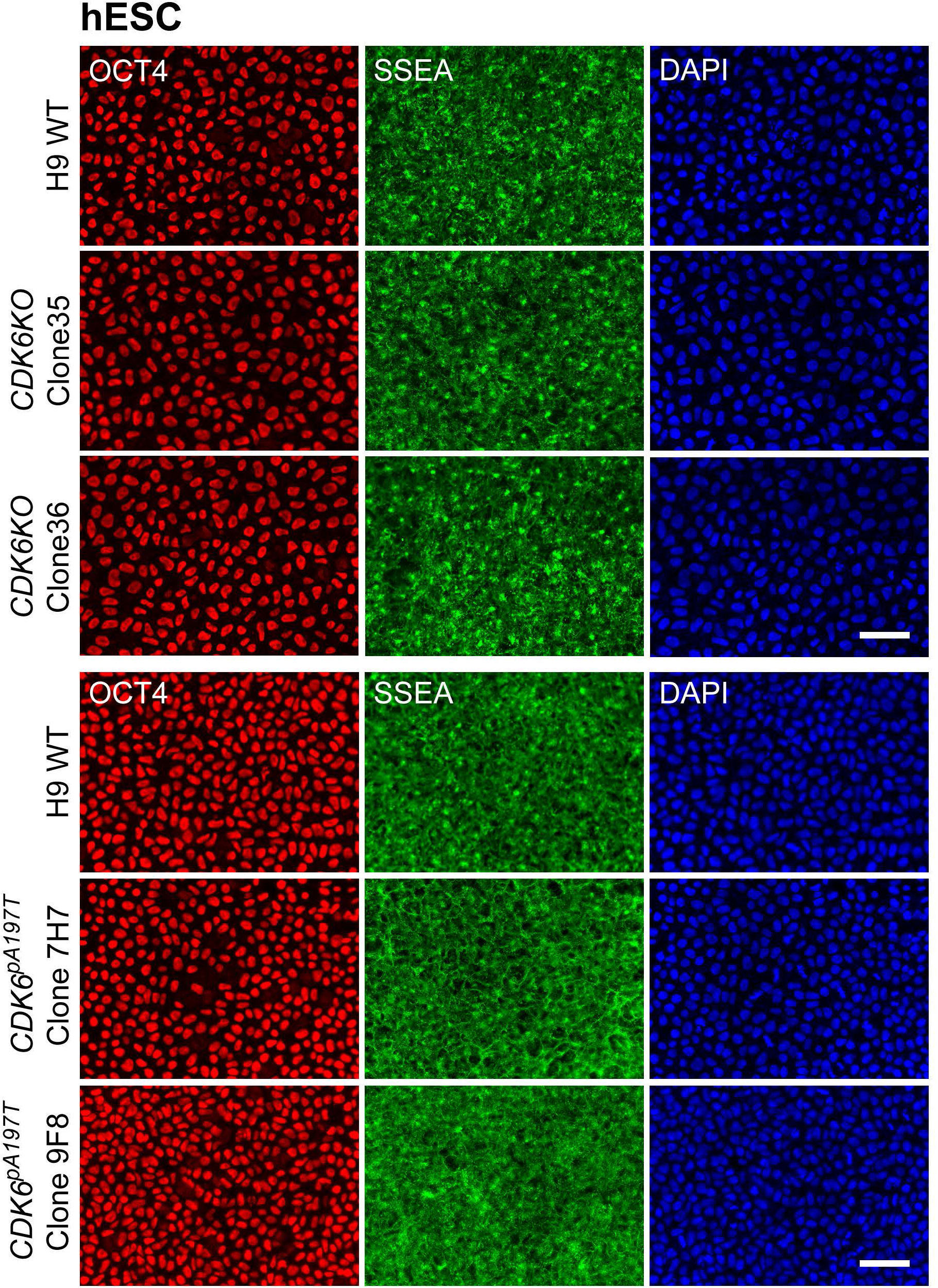
*CDK6* mutant ESCs express pluripotency markers. Micrographs of parental H9, *CDK6*^−/−^(clones 35 and 36), and *CDK6^pA197T^* (clones 9F8 and 7H7) ESCs stained for pluripotency markers OCT4 (red), SSEA (green), and DAPI (blue). Scale bar = 50 µm.

